# Rapid Peripheral Disintegration of Planaria in the Presence of Barium

**DOI:** 10.1101/2023.04.29.538822

**Authors:** Alberto Molano

## Abstract

Bioelectric signalling has been implicated in planaria regeneration and anatomical homeostasis. Previous reports demonstrated that barium chloride (1 mM), a potassium channel blocker, produced slow disintegration of the head (in 72 hours) in a clonal strain of *Dugesia japonica*. To investigate whether the effects of barium were similar in all planaria species, I tested its effects in wild *Girardia sp*. collected from a stream. The results were significantly different from those seen in *Dugesia japonica*: At 1 mM, these worms underwent complete peripheral disintegration within thirty minutes.

## Introduction

Planaria (phylum Platyhelminthes) are bilaterally symmetric, non-segmented, dorsoventrally flattened worms that possess remarkable powers of regeneration. These flatworms have a complex anatomy, with central nervous system, photoreceptors, digestive and excretory systems, epidermis, and musculature with longitudinal, circular, and diagonal fibres.

Following transverse, longitudinal, or other types of amputations, these worms can regrow the missing part and readjust their body proportions. How does the amputated fragment “know” which part is missing? For example, how does the tissue facing the anterior border of a tail fragment “know” that it is supposed to regrow a head and not a tail? Why do specific tissues regrow in the right place? How does the body “know” how to readjust its proportions?

Modern cellular and molecular biology techniques have shed light on this fascinating biological problem (1). Regeneration depends on two main factors: 1. Pluripotent stem cells called neoblasts which are spread throughout the body of the planaria, and which can differentiate into the missing cell types, and 2. Constitutive positional information, i.e., a series of “position-control genes” (PCGs) coding for receptors, ligands, or secreted inhibitors of signalling pathways with unequal concentrations along the body axes.

One example of constitutive positional information is the β-catenin protein, which is concentrated on the tails of planaria and shows a posterior-to-anterior gradient. Up- or downregulation of Wnt signalling guides whether a head or tail will regrow. Thus, inhibition of Wnt genes promotes head regeneration, whereas upregulation of this signalling pathway promotes tail formation. Similarly, Bmp signalling transmits dorsal-ventral spatial information (1). In situ hybridization and immunostaining experiments revealed that PCGs are expressed in a peripheral, subepidermal cell layer of the planaria, which turned out to be muscle cells (2).

An additional layer of complexity involved in regeneration is bioelectric signalling (3). The resting membrane potential of cells, controlled by ion channels and shared through gap junctions, seems to influence anterior-posterior polarity, and head anatomy. For example, blocking the gap junctions of amputated pre-tail fragments of *Girardia dorotocephala* for three days with octanol disturbed head regeneration, resulting in head and brain anatomies corresponding to different planaria species (4) (interestingly, the frequencies of these mismatched morphologies correlated with evolutionary distance). Similarly, perturbation of gap-junction dependent and neural signals in transversally amputated *Dugesia japonica* produced bipolar head regeneration (5).

In fact, even in non-amputated worms, blockade of ion channels can perturb their anatomy. For example, it was reported that exposure of intact *Dugesia japonica* worms to 1 mM barium chloride (BaCl_2_), a potassium channel blocker, resulted in complete head degeneration within 72 hours (6).

To investigate whether the effects of BaCl_2_ were similar in all planaria species, I tested its effects in wild *Girardia sp*. collected from a stream. Here, I report that potassium channel blockade of this particular species with barium produced significantly different effects from those reported in a clonal strain of *Dugesia japonica*. At 1 mM, these worms underwent complete peripheral disintegration within thirty minutes.

## Materials & Methods

### Planaria

Planaria were collected from Río Sáchica, a stream in Villa de Leyva, Boyacá, Colombia (5.6365° N, 73.5271° W; altitude 2,149 meters above sea level). Morphologically they appear to be *Girardia sp*. (figure 1). Worms reproduced sexually and were maintained in a tank filled with bottled water (Brisa) and oxygenated with an aquarium pump. They were fed raw beef liver once a week.

**Figure 1.**
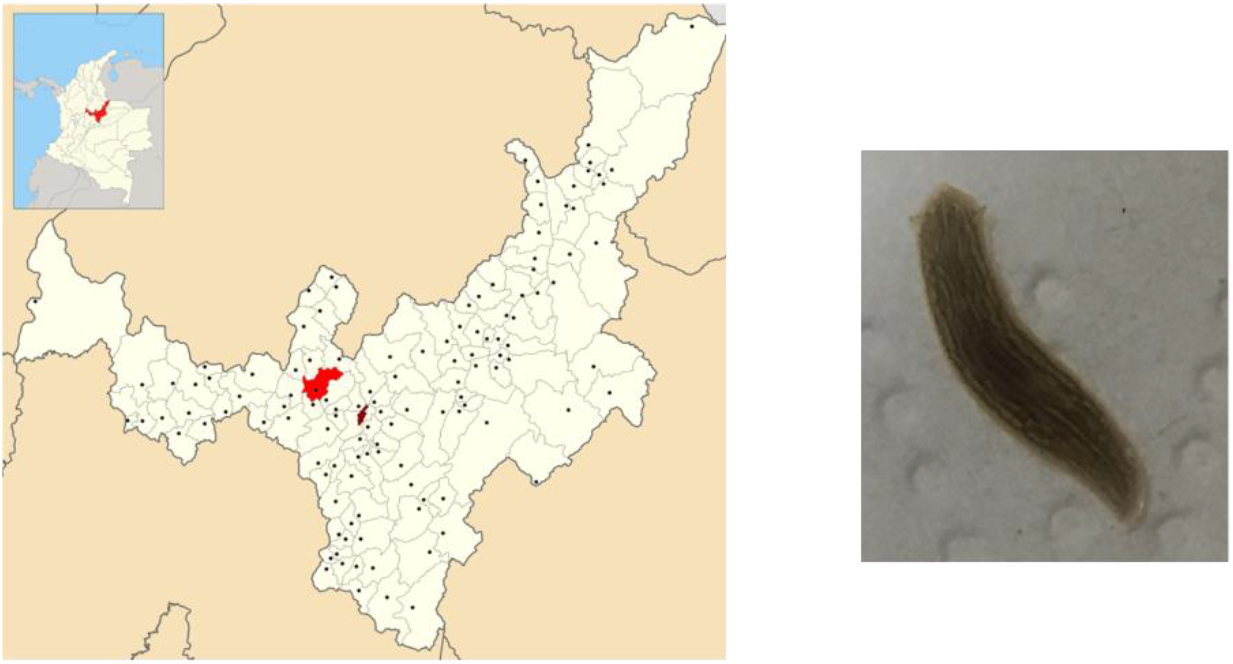
**Left:** Where the planaria were collected. Inset indicates location of the Department of Boyacá, Colombia (in red) and bigger map shows the town of Villa de Leyva. **Right:** Photograph of an adult specimen.

### Barium chloride

Barium chloride dihydrate (99% purity, batch # B401332202, calcium max. 0.05%) was purchased from Loba Chemie (Mumbai, India). Fresh 1 mM solutions were prepared in bottled water before each treatment. At this molar concentration, this solution of barium chloride dihydrate produced the characteristic repetitive “avoiding reaction” in *Paramecium* (7).

### Treatments

Freshly prepared solutions of barium chloride dihydrate (1 mM, 0.5 mM, and 0.1 mM) were added to Petri dishes. Worms were transferred with a pipette. Videos and images were captured with a smartphone and a Swift 380T microscope (Xiamen, China).

## Results

Planaria were collected from a stream in the Colombian Andes. Based on the shape of the head and auricles, as well as body pigmentation, they appear to be *Girardia sp* (figure 1). The effects of barium chloride at 1 mM were very pronounced in this planaria species (figure 2). Worms started twitching after ∼2-3 minutes. This was followed by abundant mucus secretion and pharynx extension (figure 2, 1 mM, 10 minutes). Cells and tissues started detaching from the planaria, and after just 30 minutes extensive disintegration could be seen (figure 2, 1 mM, 30 minutes; figures 3A, B, C). After one and six days all that remained was a loosely attached collection of cells (figure 2, 1 mM, 1 day, 6 days) which disaggregated with the slightest movement.

**Figure 2.**
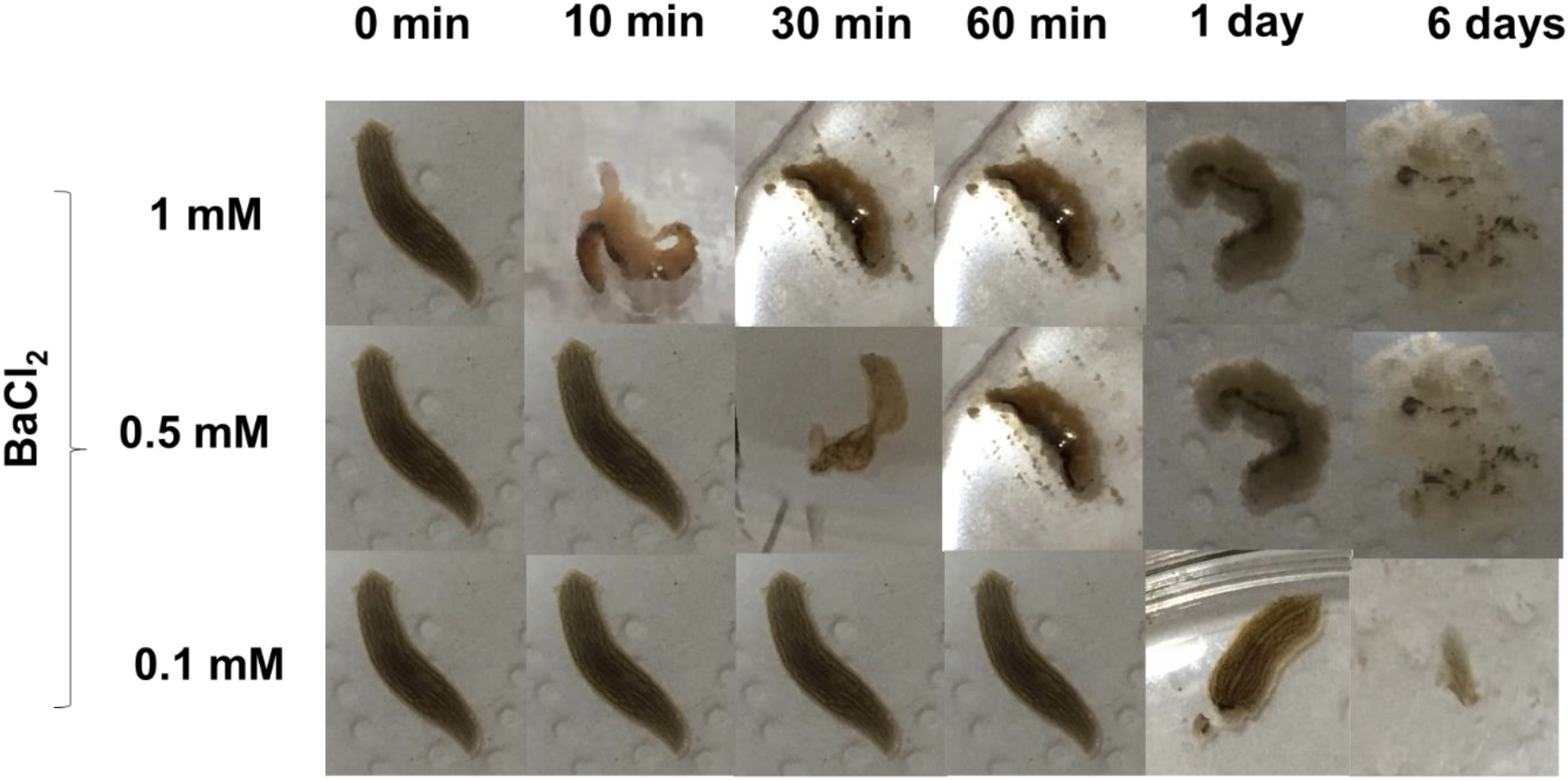
Effects of different barium concentrations (1, 0.5, and 0.1 mM) on the planaria. Images captured at 0, 10, 30 and 60 minutes, and 1 and 6 days.

**Figure 3.**
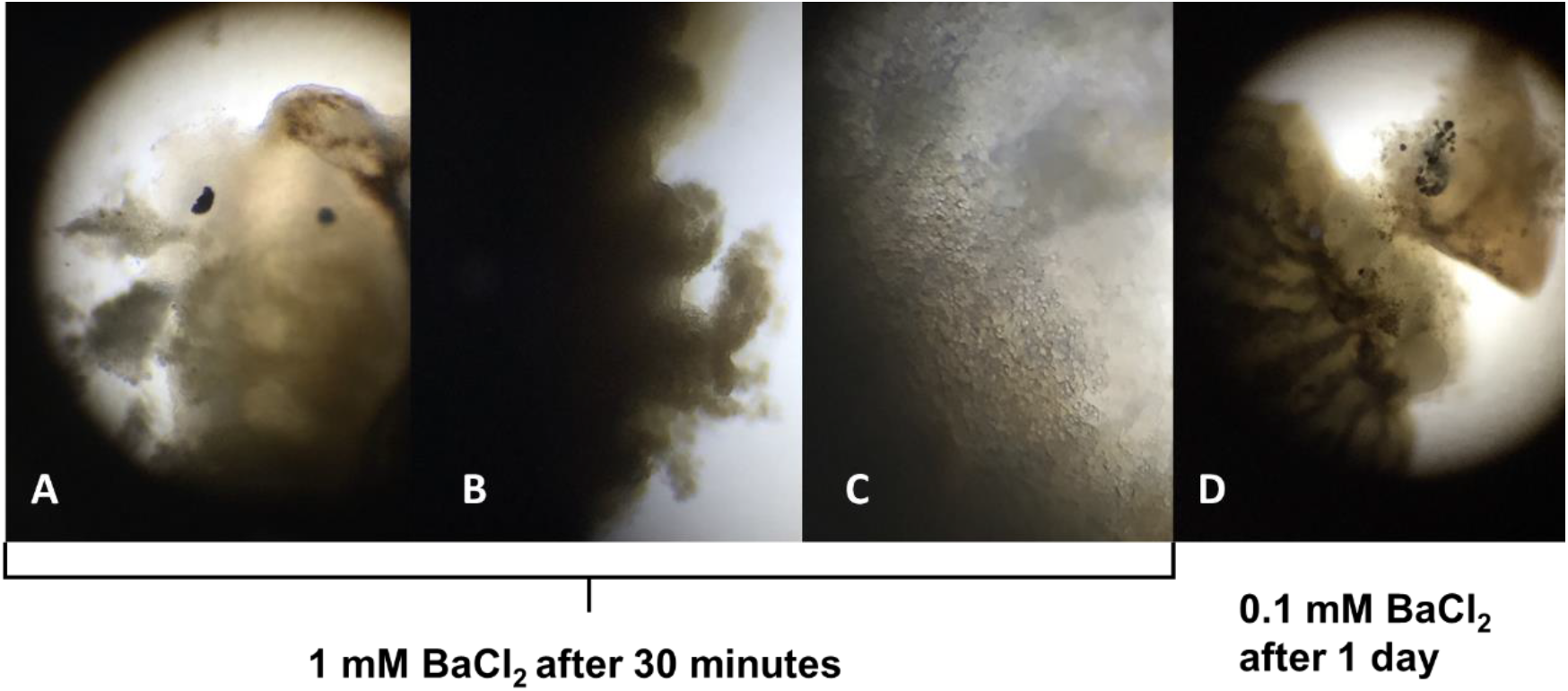
Effects of barium seen under the microscope (A, B, D: 40X; C: 100X). A, B, C: 1 mM after 30 minutes. **A**: Extensive deformation of the head, with detachment of peripheral tissue. **B, C**: Lateral detachment of tissues and cells. **D:** 0.1 mM after one day. The detached head is clearly visible.

The effects at 0.5 mM BaCl_2_ mirrored those seen at 1 mM, but with a 30-minute delay (figure 2, 0.5 mM). In contrast, the effects at 0.1 mM were only seen after 1 day, and consisted of head detachment (figure 2, 0.1 mM; figure 3D) or extrusion of the pharynx through the dorsal surface, apparently due to weakening of the musculature (not shown).

## Discussion

Previous reports of the effects of barium chloride (1 mM) on a clonal strain of *Dugesia japonica*) showed slow disintegration of the head (after 72 hours) (6). The postulated underlying mechanism involved neural excitotoxicity in the neuron-rich head of the planaria. When I tried to reproduce these experiments using wild *Girardia*, my initial expectation was that I would have to wait at least 24 hours to see anything. I was quite surprised to see the entire worms, not just the head, disintegrate in just 30 minutes.

A possible mechanism is the following: As a result of the activity of the Na^+^-K^+^-ATPase, there is a large outwardly directed K^+^ gradient in essentially all animal cells, which also show high resting membrane permeability to K^+^. As K^+^ ions diffuse out of the cell, they leave behind a negative charge inside (8). At concentrations ≥ 1 mM, barium acts as a broad-spectrum potassium channel inhibitor, preventing K^+^ ions from leaving and therefore making the inside of the cell more positive (9).

In several models, this barium-induced depolarization activates voltage-gated calcium channels. Calcium is present at very low concentrations in the cytosol and this concentration is tightly controlled since this divalent cation acts as a second messenger but can cause cell death in conditions of overload. In the published results with the clonal strain of *Dugesia japonica*, calcium channel blockers prevented barium-induced head degeneration, suggesting that Ca^++^ entry caused neurotransmitter exocytosis, particularly glutamate, subsequent neural excitotoxicity, and head degeneration (6).

However, the results presented here suggest that the effects of barium are much more widespread in other planaria species, affecting not just neurons. In other animal species like mice, barium chloride induces myofiber depolarization, opening of voltage-gated calcium channels, intracellular Ca^++^ overload, proteolysis, and membrane disruption (10). It is conceivable that a similar process is taking place in the longitudinal, circular, and diagonal muscle fibres of planaria.

Furthermore, voltage activated Ca^++^ channels play important roles not only in excitable cells but also in non-excitable ones, like ectoderm cells undergoing neural development and effector Th2 lymphocytes (11, 12). Hence, other tissues in specific planaria species, like epidermal cells, could also respond to barium by increasing their intracellular calcium concentration. This could lead to widespread activation of intracellular enzymes and cell death.

The results presented here suggest that different planaria species have different basal levels of expression of potassium channels in various tissues, and/or different sensitivities to barium blockade. This should be considered during regeneration experiments. Due to the speed with which it occurs, it is tempting to speculate that bioelectric signalling fine-tunes the minute-to-minute three-dimensional position of cells in relation to others. Future experiments will resolve these questions.

## Notes

### Competing Interest Statement

The authors have declared no competing interest.

## References

1. Reddien PW. The Cellular and Molecular Basis for Planarian Regeneration. Cell. 2018 Oct 4;175(2):327–345. doi: 10.1016/j.cell.2018.09.021. PMID: 30290140; PMCID: PMC7706840.

2. Witchley JN, Mayer M, Wagner DE, Owen JH, Reddien PW. Muscle cells provide instructions for planarian regeneration. Cell Rep. 2013 Aug 29;4(4):633–41. doi: 10.1016/j.celrep.2013.07.022. Epub 2013 Aug 15. PMID: 23954785; PMCID: PMC4101538.

3. Levin M. Molecular bioelectricity: how endogenous voltage potentials control cell behavior and instruct pattern regulation in vivo. Mol Biol Cell. 2014 Dec 1;25(24):3835–50. doi: 10.1091/mbc.E13-12-0708. PMID: 25425556; PMCID: PMC4244194.

4. Emmons-Bell M, Durant F, Hammelman J, Bessonov N, Volpert V, Morokuma J, Pinet K, Adams DS, Pietak A, Lobo D, Levin M. Gap Junctional Blockade Stochastically Induces Different Species-Specific Head Anatomies in Genetically Wild-Type Girardia dorotocephala Flatworms. Int J Mol Sci. 2015 Nov 24;16(11):27865–96. doi: 10.3390/ijms161126065. PMID: 26610482; PMCID: PMC4661923.

5. Oviedo NJ, Morokuma J, Walentek P, Kema IP, Gu MB, Ahn JM, Hwang JS, Gojobori T, Levin M. Long-range neural and gap junction protein-mediated cues control polarity during planarian regeneration. Dev Biol. 2010 Mar 1;339(1):188–99. doi: 10.1016/j.ydbio.2009.12.012. Epub 2009 Dec 21. PMID: 20026026; PMCID: PMC2823934.

6. Emmons-Bell M, Durant F, Tung A, Pietak A, Miller K, Kane A, Martyniuk CJ, Davidian D, Morokuma J, Levin M. Regenerative Adaptation to Electrochemical Perturbation in Planaria: A Molecular Analysis of Physiological Plasticity. iScience. 2019 Dec 20;22:147–165. doi: 10.1016/j.isci.2019.11.014. Epub 2019 Nov 9. PMID: 31765995; PMCID: PMC6881696.

7. Molano, A. (2023, February 2). Effects of Barium on Protozoa [Video]. YouTube. https://www.youtube.com/watch?v=boN0e3iF0h8

8. Wright SH. Generation of resting membrane potential. Adv Physiol Educ. 2004 Dec;28(1-4):139–42. doi: 10.1152/advan.00029.2004. PMID: 15545342.

9. Taglialatela M, Drewe JA, Brown AM. Barium blockade of a clonal potassium channel and its regulation by a critical pore residue. Mol Pharmacol. 1993 Jul;44(1):180–90. PMID: 8341271.

10. Morton AB, Norton CE, Jacobsen NL, Fernando CA, Cornelison DDW, Segal SS. Barium chloride injures myofibers through calcium-induced proteolysis with fragmentation of motor nerves and microvessels. Skelet Muscle. 2019 Nov 6;9(1):27. doi: 10.1186/s13395-019-0213-2. PMID: 31694693; PMCID: PMC6833148.

11. Pelletier L, Moreau M. Cav1 channels is also a story of non-excitable cells: Application to calcium signalling in two different nonrelated models. BBA - Molecular Cell Research 2021 March; 1868(6): 118996

12. Pitt GS, Matsui M, Cao C. Voltage-Gated Calcium Channels in Nonexcitable Tissues. Annu. Rev. Physiol. 2021 Feb; 83(1):183–203

